## Supplementary figures and images for "Phase separation of competing memories along the human hippocampal theta rhythm"

### Supplemental figure 1

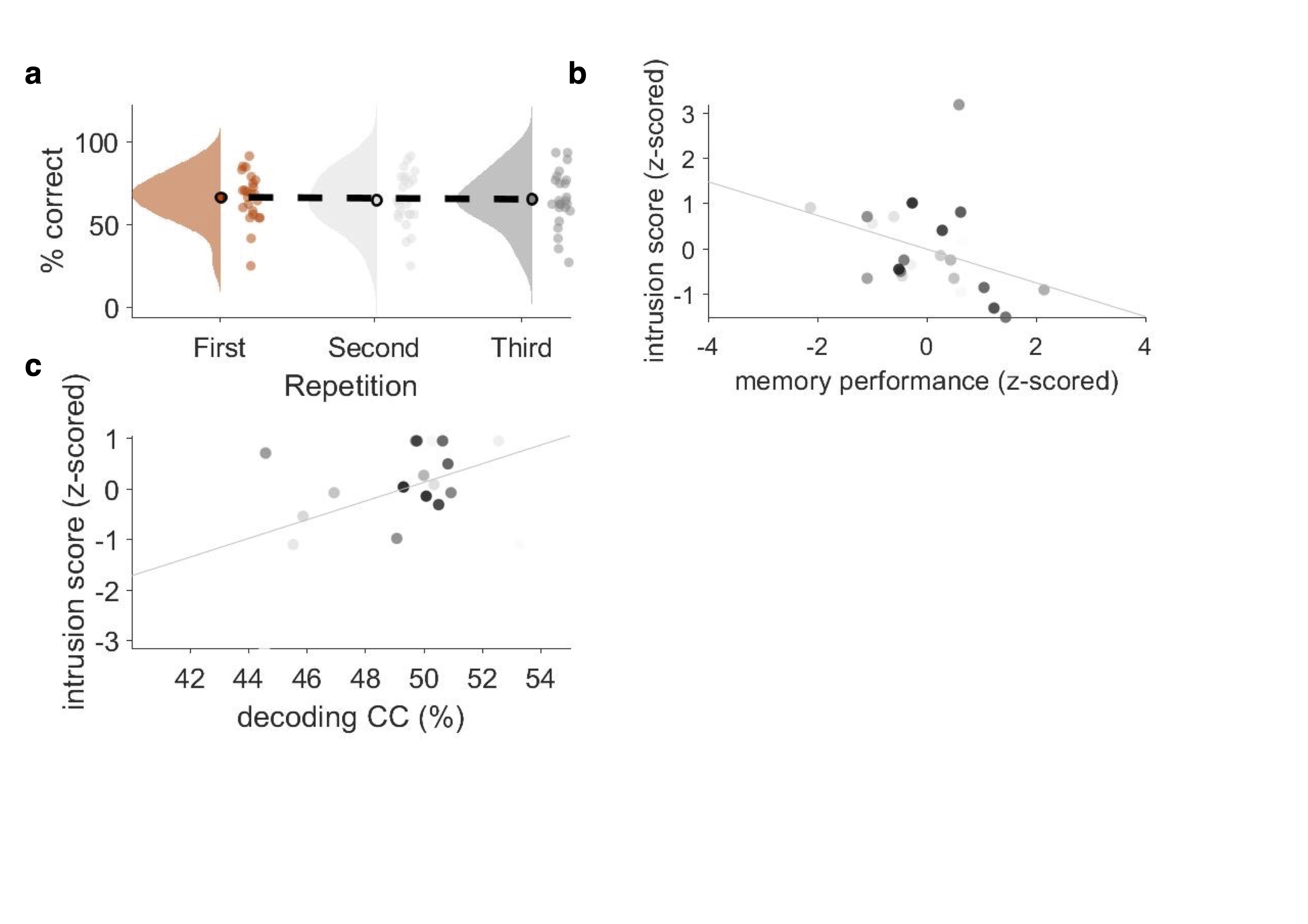

### Supplemental figure 2

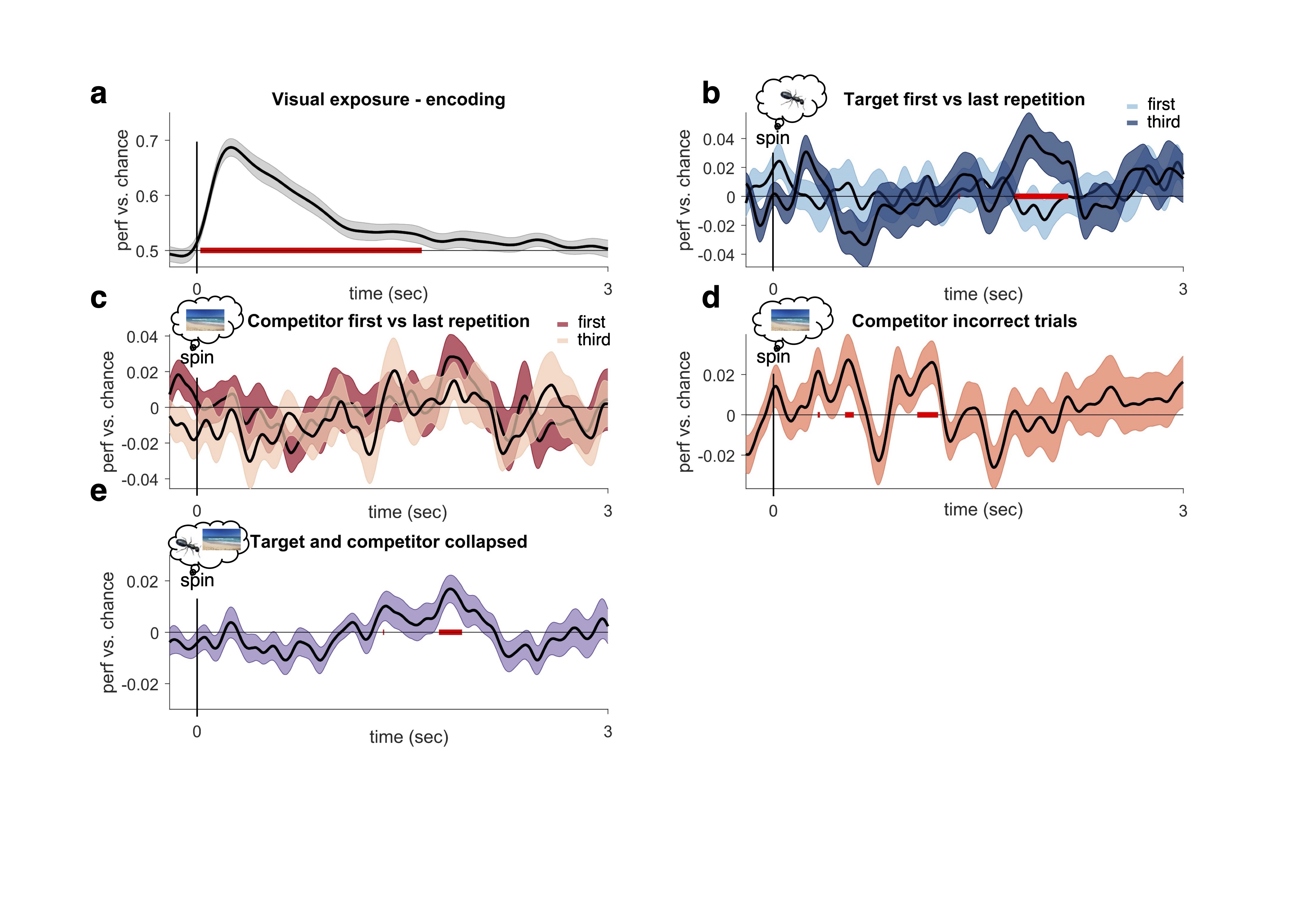

### Supplemental figure 3

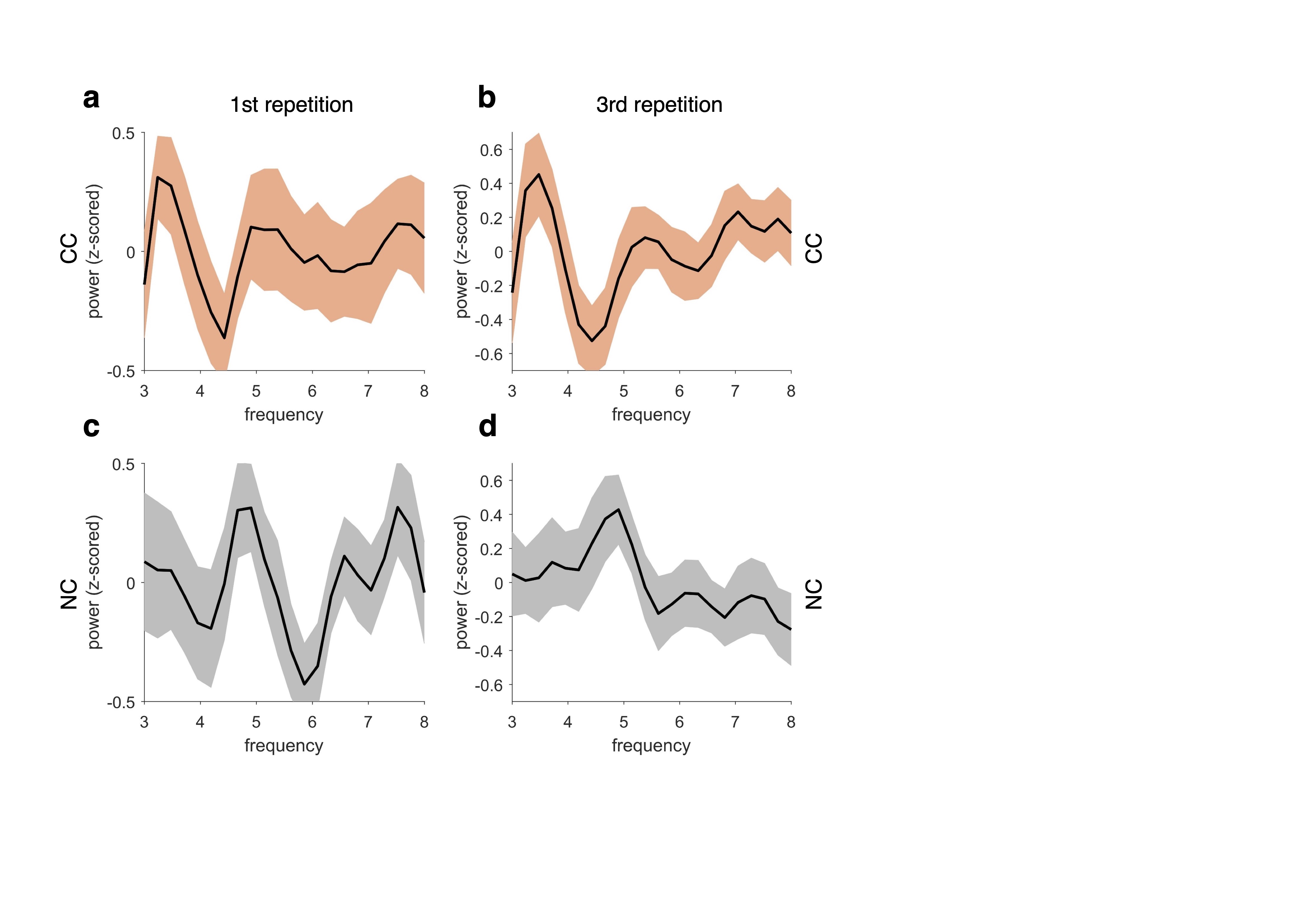

### Supplemental figure 4

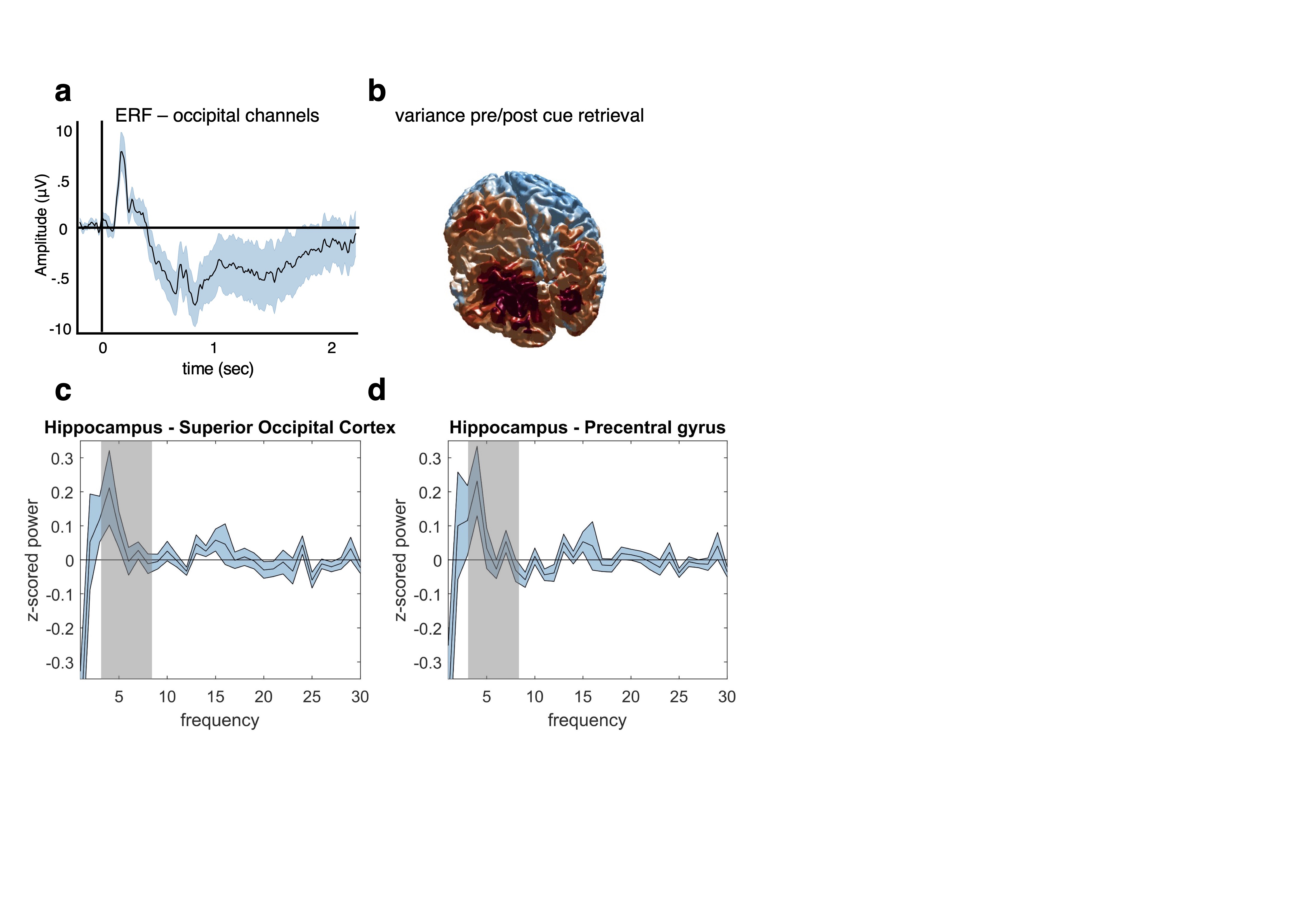

### Supplemental figure 5

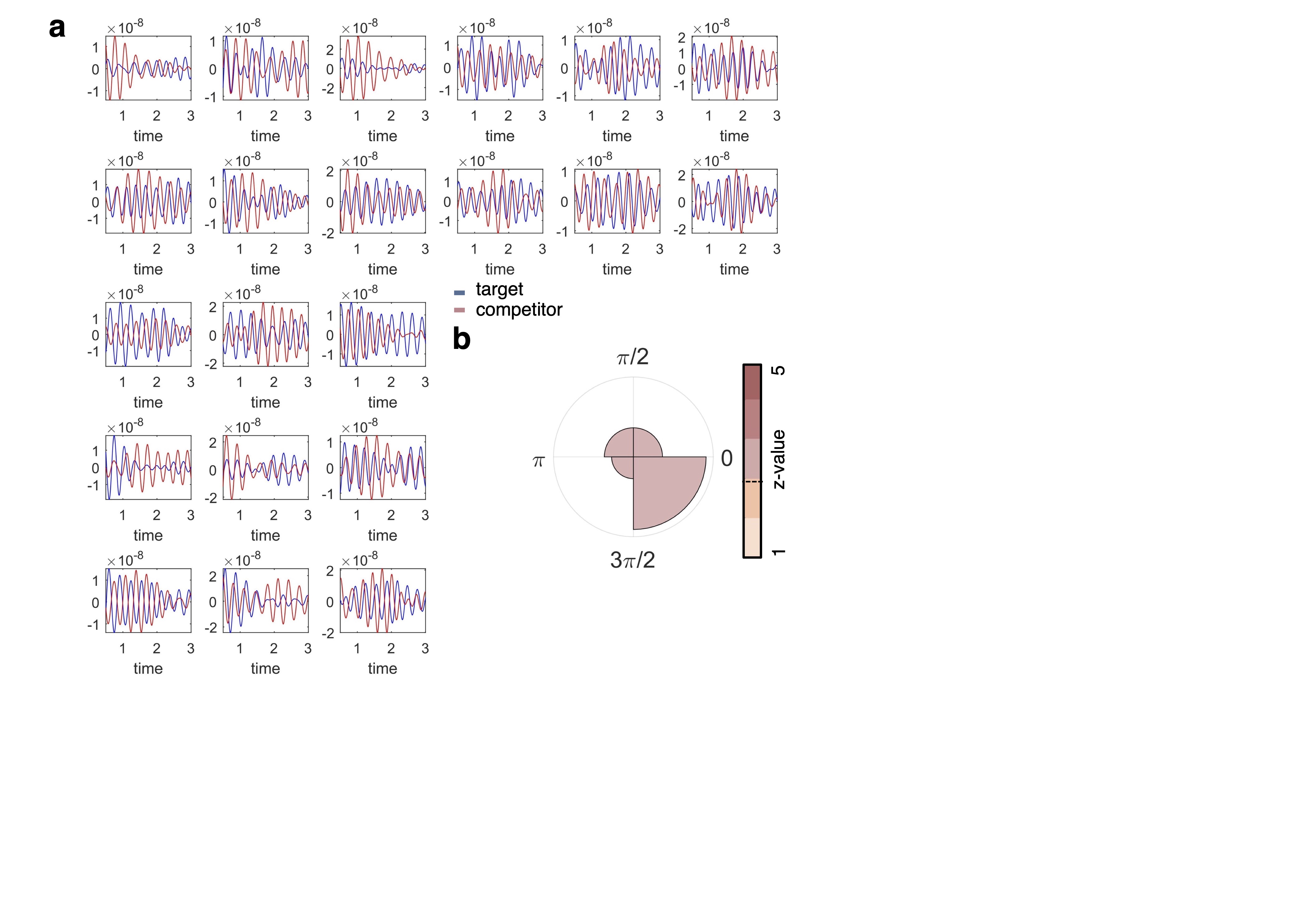
